# Toward Optimized and Predictive Connectomics at Scale

**DOI:** 10.1101/2021.01.11.426052

**Authors:** Joseph Y. Moon, Pratik Mukherjee, Ravi K. Madduri, Amy J. Markowitz, Eva M. Palacios, Geoffrey T. Manley, Peer-Timo Bremer

## Abstract

Probabilistic MRI diffusion tractography is a sophisticated technique to investigate structural connectomes, but its steep computational cost prevents application to broader research and clinical settings. Major speedup can be achieved by reducing the number of tractography streamlines. To ensure this does not degrade connectome quality, we calculate the identifiability of connectomes between test and retest MRI as a proxy for information content. We find that reducing streamline count by up to two orders of magnitude from prevailing levels in literature has no significant impact on identifiability. Incidentally, we also observe that Jaccard similarity is more effective than Pearson correlation in achieving identifiability.

*This document was prepared as an account of work sponsored by an agency of the United States government. Neither the United States government nor Lawrence Livermore National Security, LLC, nor any of their employees makes any warranty, expressed or implied, or assumes any legal liability or responsibility for the accuracy, completeness, or usefulness of any information, apparatus, product, or process disclosed, or represents that its use would not infringe privately owned rights. Reference herein to any specific commercial product, process, or service by trade name, trademark, manufacturer, or otherwise does not necessarily constitute or imply its endorsement, recommendation, or favoring by the United States government or Lawrence Livermore National Security, LLC. The views and opinions of authors expressed herein do not necessarily state or reflect those of the United States government or Lawrence Livermore National Security, LLC, and shall not be used for advertising or product endorsement purposes*.

## 1. Introduction

The structural connectome is a powerful framework for analyzing macroscale circuity of the living human brain and associating this connectivity with behavioral traits and health outcomes. Structural connectome analysis, or connectomics, has the power to distinguish autism spectrum disorder, estimate patient age and gender, and even predict cognitive ability [1, 2, 3, 4]. Furthermore, there is a significant expectation that connectomics will provide crucial insights into otherwise difficult-to-probe neurological conditions, such as traumatic brain injury (TBI) and other cognitive disorders.

However, progress in this area has been hampered by the computational cost and complexities of generating structural connectomes, particularly MRI diffusion tractography. Creating and curating connectomes for a few dozen patients, even at a research facility, may take weeks and requires dedicated personnel familiar with computational neuroscience. As a result, processing hundreds or thousands of patients for a large-scale study has been cost-prohibitive. Furthermore, computational cost remains a major impediment to developing future clinical applications that require rapid turnaround for urgent patient care needs. It is only recently that computational workflows have been developed to exploit the latest super-computers and change this paradigm [5, 6]. Here, we exploit the Department of Energy’s ability to compute large numbers of connectomes to examine the statistical stability of probabilistic tractography and the predictability of its computations. These aspects of connectomics have not been studied carefully because of the sheer scale of computational and human resources required to analyze a large number of connectomes.

In particular, we focus on probabilistic MRI diffusion tractography, by far the most resource-intensive stage of the connectomics workflow [7]. In the traditional approach, tractography can easily require thousands of CPU hours for a single subject. Traditionally, streamlines (also called fiber tracks or samples) are computed from each voxel in the white-to-grey matter boundary and connect exactly two regions of the brain. To account for the potential of crossing tracks and the uncertainty induced by the lack of spatial resolution in the MRI scans, the research standard has been to compute 1000 streamlines per boundary voxel [8]. The unofficial publication standard is as many as 5000 streamlines. However, the statistical basis for these numbers remains unclear and the results presented below suggest that there may be little to no practical benefit in computing more than 10 or 100 streamlines per voxel. This simple but significant change implies an immediate reduction in computational cost by up to two orders of magnitude without apparent loss of information.

The tractability of structural connectomes to matrix analysis has resulted in a variety of proposed techniques to associate a patient’s clinical outcomes with their connectome [9]. But since most of these techniques target specific conditions, it is difficult to use them as universal metrics for information content. In this work, we utilize a more general notion of identifiability, introduced by Amico et al. [10]. Conceptually, identifiability measures how well one can identify the connectome of a specific patient among a cohort of participants given an independently computed connectome from a prior MRI scan. The identifiability provides a generic measure of the information content of structural connectomes that is independent of any particular health condition or metric. We use a multi-center cohort of participants admitted for orthopedic, i.e. non-head related, injuries in order to demonstrate that a large streamline count does not improve identifiability in a general population. More specifically, we find that connectomes computed using 10 to 100 streamlines per voxel are as descriptive as connectomes that were generated with significantly higher streamline counts. Furthermore, the random variance induced by the probabilistic tractography is often as big as any changes observed for higher streamline counts. These two facts combined imply that many standard analyses will perform just as well with connectomes generated from a small number of streamline count than what is currently considered the standard. Reducing streamline count drastically reduces the computational resources required for the generation of structural connectomes, making structural connectomes accessible to a much wider range of researchers and paving the way for real-time connectome analysis in a clinical setting.

## 2. Methods

The tractography workflow consists of three major steps [11]: 1) calculating the probability distributions of fibers within each voxel from the raw MRI data, 2) parcellating the brain into structurally relevant regions, and 3) estimating how strongly two separate regions are connected. The main focus of this paper is to analyze heuristics for the connectivity between brain regions using different streamlines and use that information to estimate the accuracy of different levels of optimization. These heuristics must, in essence, estimate the likelihood that reconstructed connectomes match the real-world connectome. Since computing this likelihood directly is challenging, the accepted approach is to use uniform random sampling. Specifically, we begin with a large number of streamlines at each white-grey-matter boundary voxel and subsequently approximate the likelihood values by dividing the number of successful streamlines by the total number of streamlines. The likelihood values are then normalized by the volume of the regions and inserted into the connectome. Each cell of this upper-triangular matrix represents the connectivity of a region-to-region pair.

When we increase the streamline count, this process will converge to the true connectome as defined by the given parcellation and local fiber directions. As the fiber directions form a very high dimensional sampling space and a complex distribution, common wisdom would suggest that a very large number of streamlines are required for an accurate estimate. The exact origin of the accepted publication standard of streamlines, between 1000 and 5000 lines per voxel. remains unclear. But these numbers are likely the result of similar concerns regarding accuracy. However, while more streamlines undoubtedly add more information to the connectome it is not clear whether this information is relevant and/or statistically meaningful. It is well known that the physical aspects associated with an MRI procedure, i.e. measurement noise, patient motion, etc., as well as the constant change of the human brain add significant uncertainties to the measurements made on the brain which affect the generated connectome. Therefore, it is unproductive to compute the connectome to a precision that is significantly higher than the maximal resolution implied by the inherent uncertainties. However, quantitatively assessing the ‘quality’ of a connectome is not straight forward. There are two significant challenges. The first challenge is the requirement of a sufficient number of comparable MRI scans and the resources to compute their corresponding connectomes at different streamline counts. The second challenge is that there is no agreed-upon comparison metric between connectomes to understand the level of differences relevant in practice.

In this work, we address the first problem through a collaboration with the TRACK-TBI^3^ consortium [12]. TRACK-TBI is a longitudinal, observational study of TBI carried out at 18 Level 1 Trauma Centers across the United States. It includes brain-injured subjects along with a matched co-hort of orthopedic injury control subjects. All participants were followed for 12 months following injury, and MRIs were collected from a subset of both the brain-injured and orthopedic injury cohorts. To avoid potential bias from the actual brain injuries, we are using a cohort of 88 orthopedic injury control subjects all between ages 18 and 71 (mean 37.8 yr; SD 13.7 yr; 30 female). All patients have no indication of head trauma based on clinical screening. We utilize diffusion-weighted MR imaging for each patient at two time points: 2 weeks and 6 months after injury. MR imaging is conducted with 3T scanners at 11 sites across the United States. All sites use the same acquisition parameters, insofar as possible across GE, Philips, and Siemens platforms [13]. Diffusion MRI and T1-weighted MRI pre-processing and post-processing are as reported in Owen et al. [8, 14]. Given a total of 330 MRI scans we utilize a new portable and parallel computing pipeline [7] that enables us to exploit large-scale computing facilities for the necessary tractography computations.^4^ Though still under development, our pipeline accomplishes the tractography workflow using the following software components:

|  |  |  |
| --- | --- | --- |
| Segmentation | Freesurfer [15] | https://surfer.nmr.mgh.harvard.edu |
| Fiber Tensor Estimation | BEDPOSTX2 [16] | https://fsl.fmrib.ox.ac.uk |
| Probabilistic Tractography | PROBTRACKX2 [16] | https://fsl.fmrib.ox.ac.uk |

All software configurations are left to their default values, except streamline count in PROBTRACKX. Further acquisition and implementation details can be found in Moon et al. [7].

As mentioned above, in order to develop a metric to evaluate the information content in the computed connectomes, we adopted the notion of identifiability originally introduced by Amico et al. [10] in the context of functional connectomes. Information content is inherently a task-specific concept and typically used as a biomarker for various psychiatric disorders and varies accordingly. For example, the information necessary to diagnose major depressive disorder may be very different from biomarkers that indicate Alzheimer’s disease. However, despite significant advances, the analysis of structural connectomes in a clinical context remains limited and when generated, was generated in a small number of patients. As a result, there exists no widely accepted analysis of structural connectomes that could serve as a gold standard.

Identifiability provides an alternative metric based on the notion that a connectome must capture unique characteristics of the individual in order to provide any guidance towards medical outcomes or research analyses. Formulated differently, we should be able to identify a patient’s connectome within a cohort of similar patients before we expect that the connectome will be indicative of patient-specific traits. Identifiability formalizes this concept and provides a quantitative measure of how well we can identify connectomes.

The identifiability score for each patient is computed by comparing their connectome at one timepoint *p* to every connectome generated at different timepoints, *q*. As discussed in more detail below we have experimented with various forms of connectome metrics such as correlation, L2 distance, and Jaccard similarity. This results in an *N* × *N* matrix **A**, composed of correlations between the two timepoints where *N* is the number of patients. The average of diagonal elements, *I*_*self*_, measures correlation between connectomes of the same patient. The average of off-diagonals, *I*_*others*_, measures correlation between connectomes of different patients. Identifiability, *I*_*diff*_, is measured as the difference between *I*_*self*_ and *I*_*others*_.

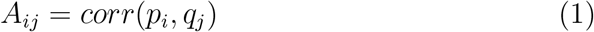

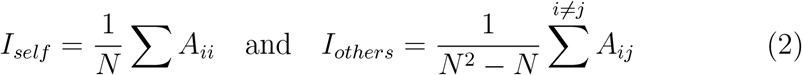

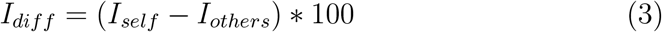

Amico and Goñi [10] improve identifiability by reducing connectome dimensionality. If we perform principal component analysis (PCA) reconstruction with *m* components, then the best possible identifiability we can extract from the data is

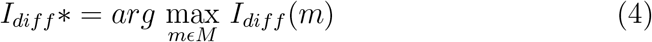

We express identifiability as equation 4 in all subsequent sections, as it represents the strongest identification ability for any set of connectomes.

Identifiability can be used to compare the success of different procedures at preserving the connectomes’ information content. However, larger study populations will necessarily have lower identifiability, since each patient must self-identify out of a wider pool of candidates. To mitigate this, we calculate the mean identifiability of repeated k-fold validation with fixed-size subsets. We randomly select a subset of *k* patients out of *n* total population, calculate identifiability of the subset, repeat this *r* times, and average the repetitions. The resulting mean identifiability enables comparison between differently sized populations.

## 3. Results

We re-ran probabilistic tractography with the same MRI scans for twenty iterations: at five streamline counts with four randoms seeds. The five streamline counts are 10, 50, 200, 500, and 1000 streamlines per voxel. In Figure 1, each data point represents the mean identifiability at a particular streamline count and random seed. Note that streamline count refers to *streamlines per voxel*. Our tractography workflow re-calculates streamlines for every region pair, so each white matter voxel at the gray-white matter boundary will actually originate many more streamlines than this number suggests [7]. We do not observe a relationship between mean identifiability and streamline count, especially considering stochastic variation and the narrow Y-axis. Since identifiability is the total percentage difference in correlation between *I*_*self*_ and *I*_*others*_ (see equation 3), small stochastic variations of fractions of a percent have little impact. However, even stochastic variation appears to have a greater impact than streamline count. This suggests that connectomes generated with low streamline counts contain just as much information as high streamline counts, at least for identification tasks.

**Figure 1:**
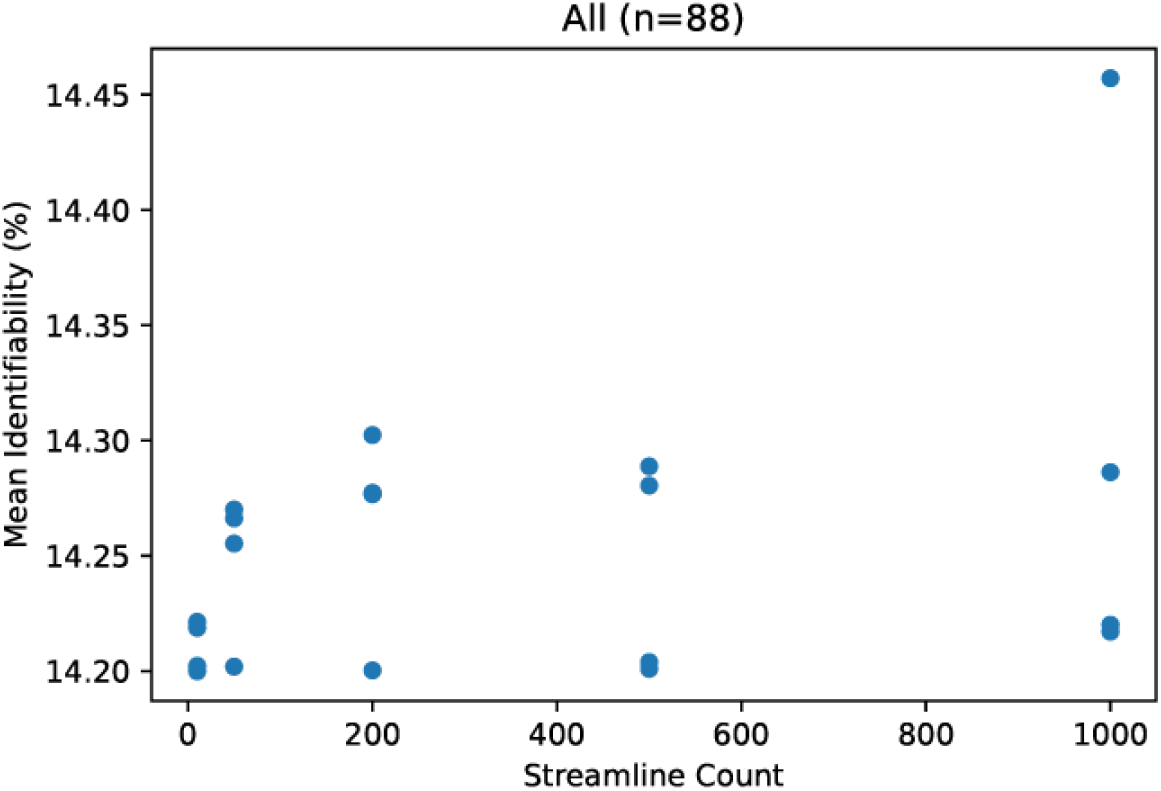
Mean identifiability with all patients (k=20, r=10)

**Figure 2:**
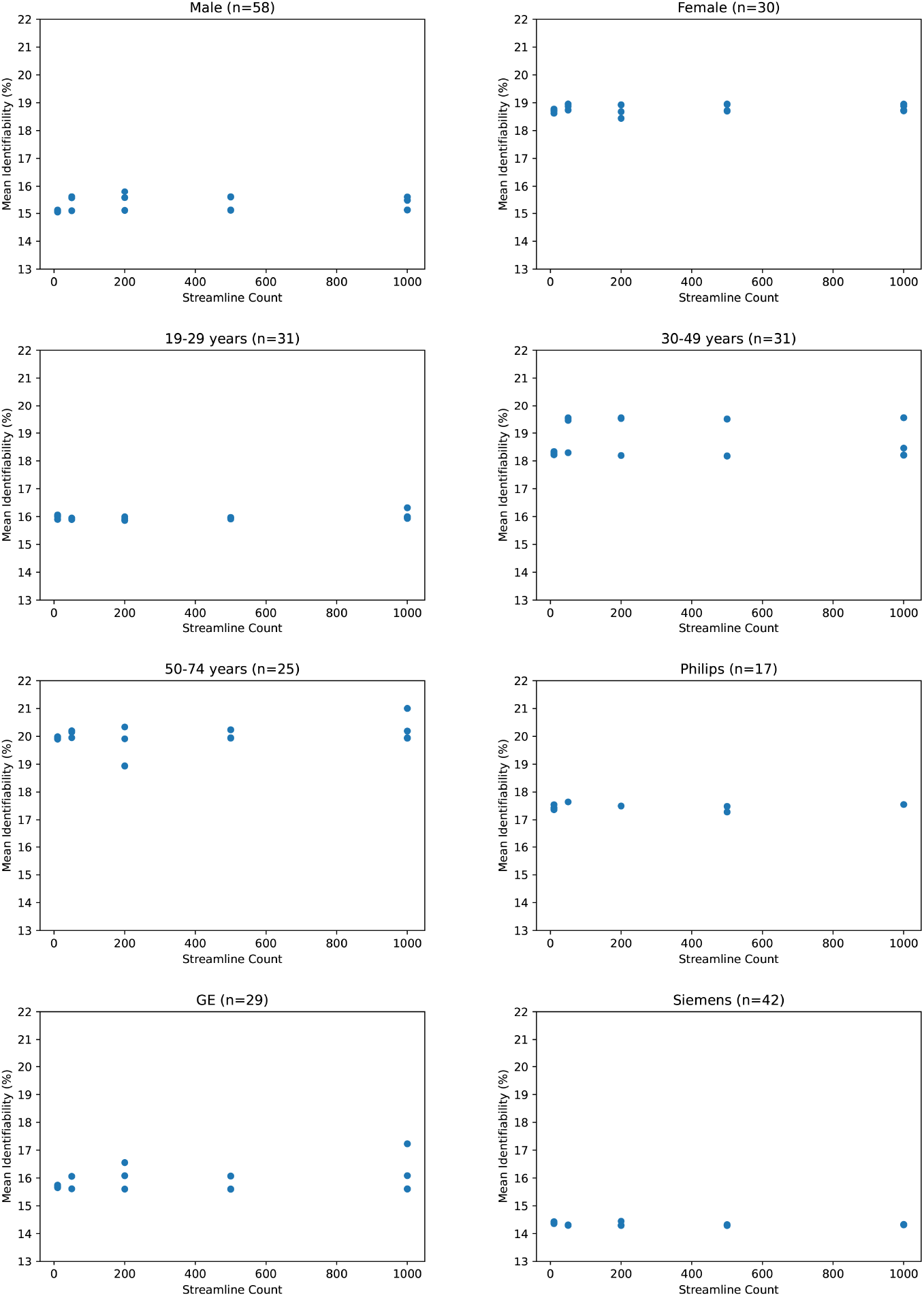
Mean identifiability by category (k=20, r=10)

If we zoom out and compare different categories, we see that mean identifiability does not have a clear positive association with streamline count no matter how patients are grouped together. We observe that certain categories present stronger differences than others. Male and female identifiability differ by 3.9 percent, the youngest and oldest patients by 4.1 percent on average, and various MRI platforms by less than 1 percent. Though this does not confirm that identifiability is reading population differences between categories, it does suggest that those differences would be more significant than any increase in identifiability from a higher streamline count.

One could argue that by comparing connectomes only against other connectomes at the same streamline count, identifiability is biased by processing artifacts unique to that streamline count. Considering this, we compared identifiability with test connectomes *p*_*i*_ against retest connectomes *q*_*j*_ from different streamline counts. Figure 3 appears to confirm this bias because identifiability is higher when the test and retest share the same streamline count. But to some degree, this is expected, as information particular to that streamline count is shared between its tests and retests, whereas those from different streamline counts may not carry that information. Nevertheless, the degree of bias does not seem to be significant compared to the overall success in identification. Again note the narrow Y-axis - even identifiability as low as 13% is more than sufficient to distinguish a retest from all 87 other retest connectomes.

**Figure 3:**
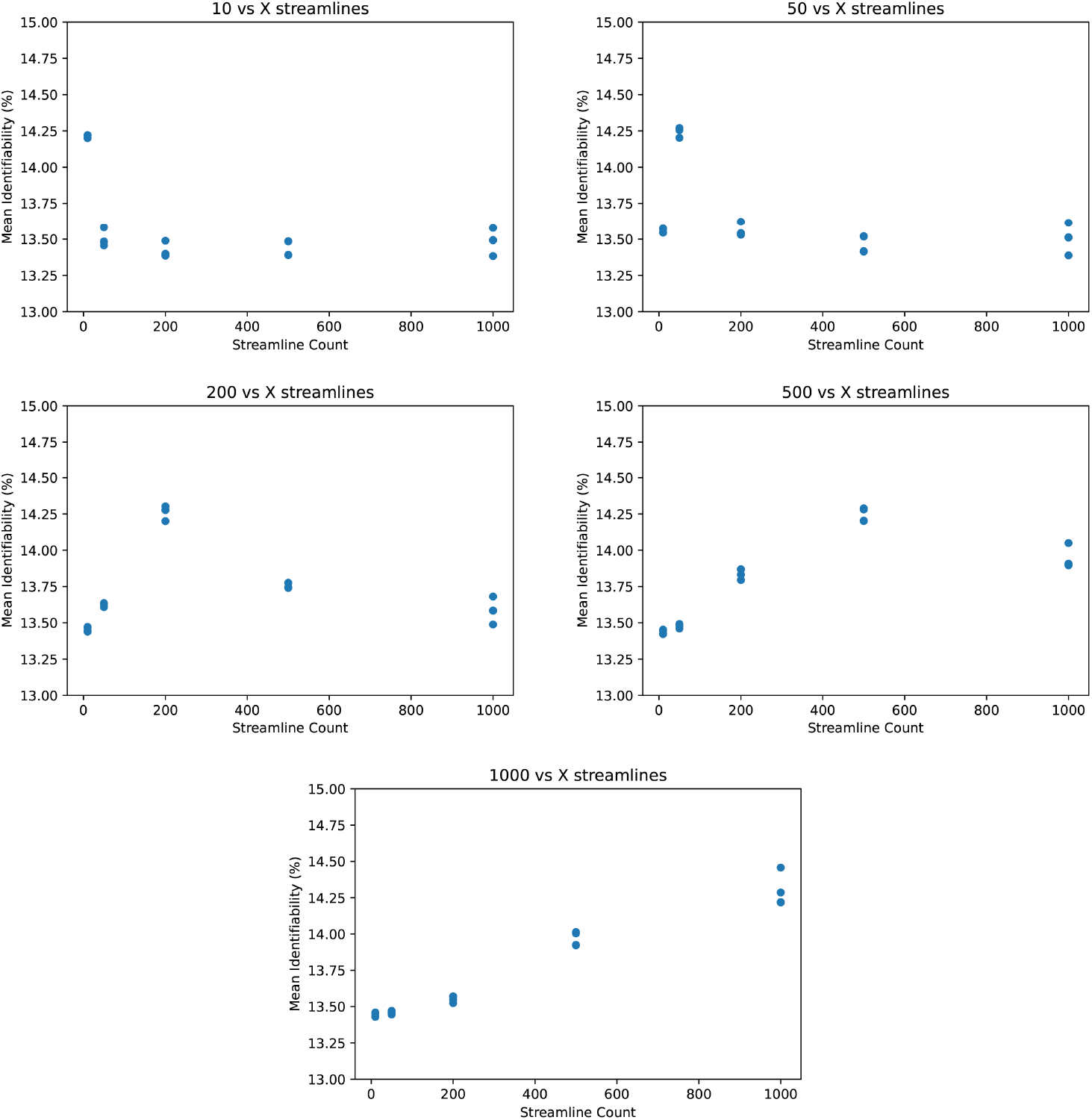
Mean identifiability across streamline counts (k=20, r=10)

Though identifiability does not encompass many of the graph analysis techniques described in the literature, it is possible to calculate identifiability using correlation metrics other than Pearson correlation. The comparison between test and retest connectomes (see equation 1) can be expressed using any linear correlation algorithm. For example, equation 5 demonstrates a comparison using L2 distance, normalized against each connectome. This metric yields somewhat better identification power than Pearson correlation. Equation 6, the normalized dot product, appears relatively weak in comparison. However, it is Equation 7, Jaccard similarity coefficient, that demonstrates significantly stronger identifiability than Pearson correlation. This is particularly unusual since Jaccard similarity discards much information from its inputs by only selecting the maximum and minimum of the test and retest values. Although we use Pearson correlation in all other figures due to its prevalence in existing literature, Figure 4 suggests that there may be room for improving the identifiability algorithm.

**Figure 4:**
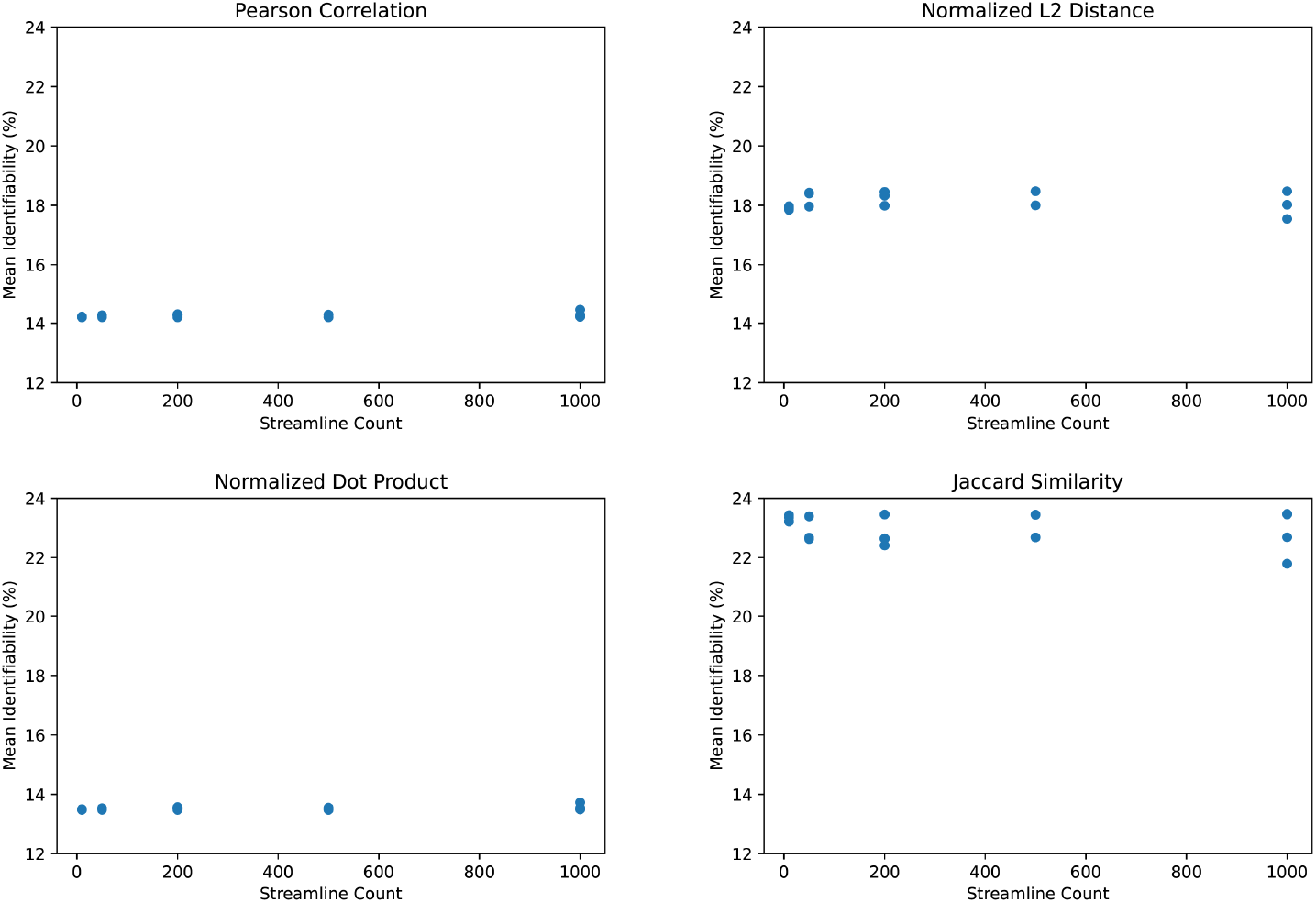
Mean identifiability by correlation metric (k=20, r=10)

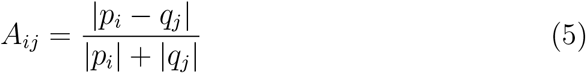

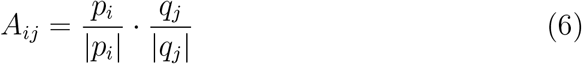

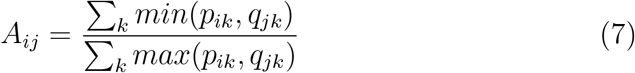

For sake of completeness, we examine the same connectomes using alternative graph metrics common in neuroimaging literature. Details of these graph metrics for the purpose of investigating test-retest reliability have been described by Owen et al. [17]. For each connectome, we 1) calculate its value using each metric and streamline count, 2) normalize by its metric value at 1000 streamlines, and 3) plot each normalized metric value in Figure 5. The resulting plots demonstrate no added value above 100 streamlines per voxel, similar to our results for identifiability.

**Figure 5:**
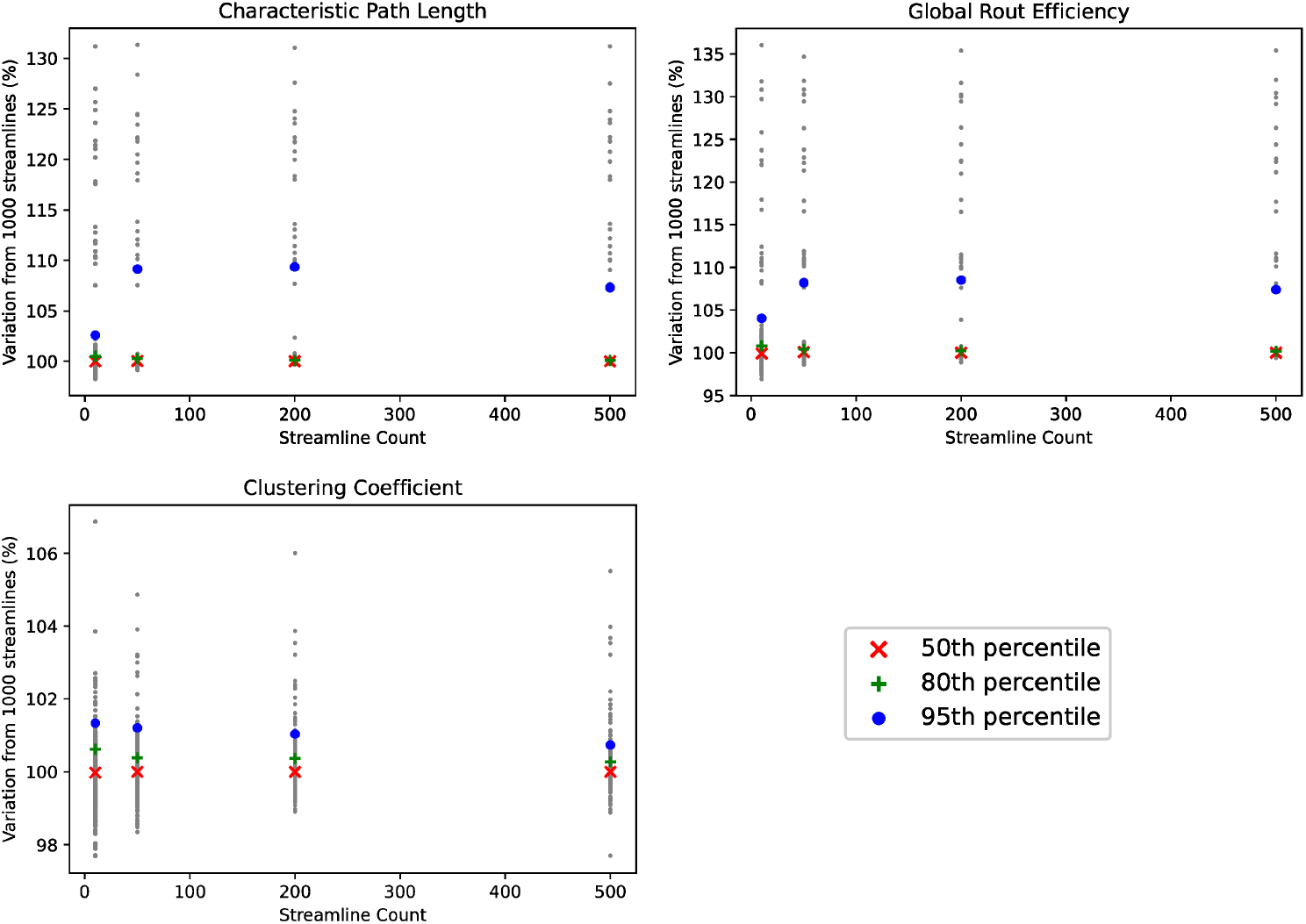
Alternative Graph Metrics

## 4. Discussion

By comparing connectomes with the concept of identifiability, we find that probabilistic tractography does not significantly benefit from high streamline counts. This has major ramifications for the computational cost and availability of tractography, as the same statistical results can be achieved with a fraction of the streamlines. However, there is a major risk that optimization would lose information not captured by identifiability. The ability to identify a patient is necessary to connectome analysis - otherwise one could argue that a connectome is indistinguishable and therefore dominated by noise and external variables. But even if we could perfectly identify patients from connectomes, this may not be sufficient for more complex analyses.

There is also the risk that we did not compute sufficient samples. To address this, we re-ran probabilistic tractography on all patients with five streamline counts and four different random seeds, for a total of twenty iterations. With that amount of data, streamline count and identifiability do not appear to be correlated. However, it is remotely possible that running far more than twenty iterations would show correlation instead. We do not pursue this possibility owing to the computational expense of tractography with high streamline counts - generating our data already consumed over 300,000 CPU hours.

Another limitation of this work is the use of a specific probabilistic streamline tractography approach using a specific method of gray matter atlas parcellation for connectome reconstruction. These results will need to be reproduced and generalized across other tractography methods and other techniques for gray matter parcellation. There is also the concern that these findings lack external physiological data. Brains do not exist in a vacuum, so key markers such as clinical survey results, blood pressure, and body weight may influence connectome analysis in subtle ways. We mitigate this to an extent by categorizing patients by age and gender and find that nothing in these categories undermines our argument regarding streamline count. However, we do not possess further physiological data for this patient population, so the influence of other external variables remains unexplored.

We also find that Jaccard similarity outperforms more commonly used connectome correlation metrics such as Pearson correlation in the calculation of identifiability. Though we are surprised that this is the case, it is possible that Jaccard similarity increases the weight of low-frequency information by effectively binarizing the non-shared values. When calculating identifiability, high-frequency values, such as dense contiguous sections of the brain, may often match to the wrong subject. Subjects are better distinguished by low-frequency areas with unique structures. Given an incorrect match, choosing a minimum or maximum of the test and retest value in low-frequency areas will create a strongly fluctuating test-retest variation since values tend not to overlap. And whereas Pearson correlation and other metrics would dilute this variation by the weight of high-frequency areas, Jaccard similarity would provide consistent test-retest variation in high-frequency areas since it does not combine the test and retest in each voxel. As a result, Jaccard similarity improves identifiability similarly to PCA reconstruction, by pruning low-information data. However, this is mostly speculation and would require further study beyond the scope of this paper.

## 5. Conclusions

Progress in connectomics has been limited by the steep computational cost of probabilistic white matter fiber tractography. Creating diverse datasets with large numbers of patients requires optimizations of the tractography workflow. However, excessive optimization may degrade the connectome’s information content. To measure the extent to which we can optimize tractography, we use identifiability as an approximate measure of the average information content in a set of connectomes. Identifiability is a quantifiable metric for identification tasks predictiveness using a patient’s test and retest, based on MRIs conducted six months apart. This enables us to optimize computation right up to the threshold of information loss.

Probabilistic tractography is computationally expensive because it simulates massive quantities of white-matter fiber streamlines. We find that the number of streamlines can be greatly reduced from current practice. This optimization appears to have no impact on identifiability; ergo, it does not degrade the connectome’s information content for most purposes. Reducing the number of streamlines yields direct linear efficiencies, such that using half the streamlines takes approximately half the time to compute. Existing literature uses between 1000 and 5000 streamlines per voxel to ensure a well-converged solution. We find that identifiability is stable with as few as 10 streamlines per voxel.

We find that low streamline counts perform just as well as high streamline counts even when analyzing our study population with different demographics. These findings hold true for male and female patients, different age ranges, different correlation metrics, and all three common MRI hardware platforms. The choice of population makes a far greater impact than any decision on streamline count. In fact, variations in mean identifiability due to streamline count are even less than those from stochastic variation due to probabilistic tractography.

Using low streamline counts promises to greatly accelerate connectome analysis. High streamline counts do not appear to harm identifiability in any scenario, and will likely continue to be the standard for small-scale studies. But by reducing the computational cost of tractography, this simple optimization will enable hundreds to thousands of connectomes to be generated on systems that previously handled a few dozen. Many open neuroimaging questions cannot be answered with small-scale studies alone, particularly those related to subtle population differences such as behavioral disorders. As the field of connectomics grows, optimizations such as these will be necessary to keep up with the large amount of clinical data and computational resources applied to human brain research as well as foster clinical applications that require faster results for real-time patient care.

## Additional TRACK-TBI Investigators

Opeolu Adeoye, MD; Neeraj Badjatia, MD; Kim Boase; Jason Barber, MS; Yelena Bodien, PhD; M. Ross Bullock, MD PhD; Randall Chesnut, MD; John D. Corrigan, PhD; Karen Crawford,; Ramon Diaz-Arrastia, MD PhD; Sureyya Dikmen, PhD; AnnChristine Duhaime, MD; Richard Ellenbogen, MD; V Ramana Feeser, MD; Adam R. Ferguson, PhD; Brandon Foreman, MD; Raquel Gardner; Etienne Gaudette, PhD; Joseph Giacino, PhD; Dana Goldman, PhD; Luis Gonzalez; Shankar Gopinath, MD; Rao Gullapalli, PhD; J Claude Hemphill, MD; Gillian Hotz, PhD; Sonia Jain, PhD; C. Dirk Keene, MD PhD; Frederick K. Korley, MD; Joel Kramer, PsyD; Natalie Kreitzer, MD; Harvey Levin, MD; Chris Lindsell, PhD; Joan Machamer, MA; Christopher Madden, MD; Alastair Martin, PhD; Thomas McAllister, MD; Michael McCrea, PhD; Randall Merchant, PhD; Lindsay Nelson, PhD; Laura B. Ngwenya, MD; Florence Noel, PhD; Amber Nolan, MD PhD; David Okonkwo, MD PhD; Daniel Perl, MD; Ava Puccio, PhD; Miri Rabinowitz, PhD; Claudia Robertson, MD; Jonathan Rosand, MD; Angelle Sander, PhD; Gabriella Satris; David Schnyer, PhD; Seth Seabury, PhD; Sabrina Taylor, PhD; Nancy Temkin, PhD; Arthur Toga, PhD; Alex Valadka, MD; Mary Vassar, RN MS; Paul Vespa, MD; Kevin Wang, PhD; John K. Yue, MD; Esther Yuh, MD PhD; Ross Zafonte.

## Affiliations of Additional TRACK-TBI Investigators: University

of Cincinnati (Adeoye, Foreman, Kreitzer, Ngwenya); University of Maryland (Badjatia, Gullapalli); University of Washington (Barber, Boase, Bodien, Chesnu, Dikmen, Ellenbogen, Keene, Machamer, Temkin); Massachusetts General Hospital (Bodien, Rosand); University of Miami (Bullock, Hotz); Ohio State University (Corrigan); University of Southern California (Crawford, Gaudette, Goldman, Seabury, Toga); University of Pennsylvania (Diaz-Arrastia); Massachusetts General Hospital for Children (Duhaime); Virginia Commonwealth University (Feeser, Merchant, Valadka); University of California, San Francisco (Ferguson, Gardner, Hemphill, Kramer, Martin, Nolan, Satris, Taylor, Vassar, Yue, Yuh); Spaulding Rehabilitation Hospital (Giacino); TIRR Memorial Hermann (Gonzalez); Baylor College of Medicine (Gopinath, Levin, Noel, Robertson, Sander); University of California, San Diego (Jain); University of Michigan (Korley); Vanderbilt University (Lindsell); University of Texas Southwestern Medical Center (Madden); Indiana University (McAllister); Medical College of Wisconsin (McCrea, Nelson); Uniformed Services University (Perl); University of Pittsburgh (Okonkwo, Puccio, Rabinowitz); University of Texas, Austin (Schnyer); University of California, Los Angeles (Vespa); University of Florida (Wang); Harvard Medical School (Zafonte).

## Acknowledgements

The research is funded by the United States Department of Energy under the DOE Office of Science, Advanced Scientific Computing Research. Support is organized under The Co-Design for Artificial Intelligence and Computing at Scale for Extremely Large, Complex Datasets projects (Grant #KJ040301).

Geoffrey Manley discloses grants from the United States Department of Defense – TBI Endpoints Development Initiative (Grant #W81XWH-14-2-0176), TRACK-TBI Precision Medicine (Grant #W81XWH-18-2-0042), and TRACK-TBI NETWORK (Grant #W81XWH-15-9-0001); NIH-NINDS – TRACK-TBI (Grant #U01NS086090); and the National Football League (NFL) Scientific Advisory Board – TRACK-TBI LONGITUDINAL.

The United States Department of Energy supports Dr. Manley for a precision medicine collaboration. One Mind has provided funding for TRACK-TBI patients stipends and support to clinical sites. He has received an unrestricted gift from the NFL to the UCSF Foundation to support research efforts of the TRACK-TBI NETWORK. Dr. Manley has also received funding from NeuroTruama Sciences LLC to support TRACK-TBI data curation efforts. Additionally, Abbott Laboratories has provided funding for add-in TRACK-TBI clinical studies.

Amy Markowitz receives funding from the Department of Defense TBI Endpoints Development Initiative (Grant #W81XWH-14-2-0176) and TRACK-TBI NETWORK (Grant #W81XWH-15-9-0001). Ms. Markowitz also receives salary support from the United States Department of Energy precision medicine collaboration and the philanthropic organization, One Mind.

## Footnotes

3 Transforming Research and Clinical Knowledge in Traumatic Brain Injury (https://tracktbi.ucsf.edu)

4 Publicly available at https://github.com/LLNL/MaPPeRTrac

## Notes

### Competing Interest Statement

The authors have declared no competing interest.

